# GALR2 W248L mutation exacerbates neuroinflammation through pro-inflammatory macrophage polarization and microglial activation in experimental autoimmune encephalomyelitis

**DOI:** 10.64898/2026.07.27.740554

**Authors:** Raphael Morales-Neto, Danieli Cristina Gonçalves, Karina Yumi Degaki, João Paulo Mesquita Luiz, Luis Eduardo Alves Damasceno, Matheus Severino Brandemarte, Santiago José Ortiz-Peñuela, André Almeida Schenka, Emmanuel Dias-Neto, Alexandre Leite Rodrigues de Oliveira, José Carlos Alves-Filho, Daniela Barretto Barbosa Trivella, Ângela Saito

**Affiliations:** Brazilian Center for Research in Energy and Materials (CNPEM), Brazilian Biosciences National Laboratory (LNBio), Campinas, Brazil; Graduate Program in Molecular and Morphofunctional Biology (PPG-BMM), University of Campinas (UNICAMP), Campinas, Brazil; Department of Structural and functional Biology, Instituto de Biologia, Universidade de Campinas (UNICAMP), Campinas, Brazil; Department of Pharmacology, Ribeirão Preto Medical School, University of Sao Paulo, Ribeirão Preto, Brazil; Center for Research in Inflammatory Diseases, Ribeirão Preto Medical School, University of Sao Paulo, Ribeirão Preto, Brazil; Faculty of Medical Sciences, Department of Pharmacology, University of Campinas (UNICAMP), Campinas, Brazil; Rutgers Cancer Institute and Division of Cancer Biology, Department of Radiation Oncology, Rutgers New Jersey Medical School, Newark, New Jersey, USA

**Keywords:** CRISPR/Cas9, GALR2, EAE, microglia, inflammasome, multiple sclerosis, neuroinflammation

## Abstract

Multiple sclerosis (MS) is a chronic neuroinflammatory disease characterized by demyelination, neurodegeneration, and progressive neurological disability. Galanin, a neuropeptide with immunomodulatory properties, signals through G protein-coupled receptors, among which galanin receptor 2 (GALR2) has been implicated with neuroprotective and anti-inflammatory functions. A rare homozygous single nucleotide variant in *GALR2* (rs61745847; p.W249L) has been identified in a patient diagnosed with relapsing-remitting MS, however, the biological relevance of this variant in neuroinflammation remains unknown. Here, we investigated the impact of the orthologous *GALR2* W248L mutation using a knock-in mouse model and experimental autoimmune encephalomyelitis (EAE). GALR2 W248L knock-in (KI) mice exhibited a more severe clinical course of EAE, accompanied by enhanced inflammatory infiltration, exacerbated demyelination, and increased microglial activation in the spinal cord compared with wild-type (WT) mice. Despite comparable lymphoid and myeloid cell frequencies in the central nervous system, alterations in microglial density and morphology suggested an important contribution of the innate immune system to disease exacerbation in the KI mice. *Ex vivo* analyses revealed that bone marrow-derived macrophages from KI animals exhibited a pronounced shift toward a pro-inflammatory phenotype, characterized by enhanced M1 polarization, impaired M2-associated responses, and increased NLRP3 inflammasome activation. In parallel, live-cell imaging of primary hippocampal neurons demonstrated reduced galanin binding in mutant cells, consistent with impaired GALR2 functional availability at the plasma membrane. Together, these findings identify GALR2 as a modulator of the neuroinflammatory response and indicate that disruption of galanin-GALR2 signaling promotes sustained innate immune activation, highlighting the relevance of this pathway for MS pathogenesis and its potential as a therapeutic target in neuroinflammatory disorders.

## Introduction

Multiple sclerosis (MS) is a chronic, inflammatory disease of the central nervous system (CNS), characterized by demyelination, neurodegeneration, and progressive neurological impairment. Its etiology is multifactorial, arising from a complex interplay between genetic susceptibility and environmental factors [1,2]. Although MS has traditionally been viewed as a T cell–mediated autoimmune disorder, growing evidence underscores the contribution of B cells, microglia, and other components of the innate immune system to disease pathogenesis. Clinically, MS presents in various forms, most commonly beginning with relapsing-remitting episodes that may transition into secondary progressive disease. MS affects an estimated 2.9 million people worldwide, with an average global incidence of 2.1 cases per 100,000 population per year, equivalent to roughly one new diagnosis every 5 minutes, making it a leading cause of non-traumatic disability among young adults worldwide. [1,2].

The main pathological hallmark of MS is the presence of demyelinating lesions in the brain and spinal cord, driven by immune cell infiltration following blood-brain barrier (BBB) disruption [1]. These lesions derive from Infiltrating immune cells, together with activated resident microglia and astrocytes, after the release of pro-inflammatory mediators that contribute to demyelination and neuro-axonal damage [1,3,4]. While adaptive immune mechanisms have long been the primary focus of MS research, increasing attention has been directed toward innate immune cells, such as microglia, macrophages, and dendritic cells. These cells not only respond to damage-associated molecular patterns (DAMPs) but also activate NLRP3 inflammasomes and amplify inflammation within the CNS [4,5,6]. The NLRP3 inflammasome functions as a key amplifier of inflammatory responses through the maturation of interleukin-1β (IL-1β) [7,8]. IL-1β acts as a pivotal effector in MS and experimental autoimmune encephalomyelitis (EAE), promoting leukocyte recruitment, BBB disruption, and glial activation, thereby exacerbating neuroinflammation and demyelination [8,9]. Consistently, while increased IL-1β levels and NLRP3 activity are observed in active MS lesions, the targeting of NLRP3 signaling attenuates disease severity in EAE models [8,10,11,12]. Collectively, these findings identify IL-1β as a central effector linking innate immune activation to sustained neuroinflammation in demyelinating diseases.

Among the neuropeptides implicated in immune modulation, galanin has emerged as a relevant player in neuroinflammatory contexts [13]. It mediates its effects through three G protein-coupled receptors (GPCRs), namely GALR1, GALR2, and GALR3 [8,9]. Of these, GALR2 displays the most versatile signaling capacity, coupling to both G_q/11_ and G_i/o_ proteins, thus modulating pathways such as phospholipase C/protein kinase C (PLC/PKC) and cyclic AMP/protein kinase A (cAMP/PKA), respectively [15,16,17,18]. In addition to its expression in the hippocampus and other regions of the CNS, GALR2 is also expressed in immune cells, including T cells, B cells, and macrophages, highlighting its dual role at the interface between the nervous and immune systems [17,19, 20,21]. Consistent with this expression pattern, GALR2 has been associated with neuroprotective, anti-inflammatory, and axonal growth-promoting functions [20–22].

Accumulating evidence implicates the galanin-GALR2 axis as a critical modulator of immune responses and glial activation in inflammatory diseases, including MS. Notably, galanin expression is elevated in active MS lesions, and genetic deletion of galanin or GALR2 (knock-out) in mice exacerbates inflammation and neurodegeneration in experimental autoimmune encephalomyelitis (EAE) model, the most widely used murine model to study MS-like pathology [23]. Collectively, these observations support the hypothesis that intact GALR2 signaling contributes to immune regulation and neuroprotection, and that its disruption may promote disease severity.

In this context, a rare homozygous single-nucleotide variant (SNV; rs61745847) in the human *GALR2* gene, resulting in a tryptophan-to-leucin substitution (GALR2 W249L), was identified in a Brazilian patient with sporadic relapsing-remitting MS, after whole exome sequencing of a cohort of patients diagnosed with sporadic MS and their biological parents [24]. Functional studies indicate that this variant impairs galanin signaling by reducing GALR2 availability at the plasma membrane, thereby disrupting downstream signal transduction [24]. To investigate the biological relevance of this mutation *in vivo*, we generated a knock-in mouse line carrying the orthologous W248L mutation in *Galr2* using CRISPR/Cas9-mediated genome editing. Using the EAE model, we examined the impact of impaired GALR2 signaling on neuroinflammatory responses by assessing disease progression, histopathological alterations, glial activation, cytokine production, and immune cell profiles in homozygous knock-in (KI) and wild-type (WT) mice. In parallel, we performed *ex vivo* assays to evaluate galanin receptor binding in primary neurons, as well as macrophage polarization and inflammatory responsiveness in bone marrow-derived macrophages from naïve WT and KI animals. Together, our integrated *in vivo* and *ex vivo* approach indicates that the GALR2 W248L mutation impairs galanin signaling and implicates the galanin-Galr2 axis in the neuroinflammatory processes that govern disease severity in this model of MS-like pathology.

## Materials and methods

### Generation of GALR2 W248L Knock-in Mice and Genotyping

The GALR2 W248L (c.743G>C p.S248W>L) mouse line was generated through CRISPR/Cas9 genome editing tool by the Genome Editing Laboratory (LEG) at LNBio/CNPEM. The single guide RNA (sgRNA) was generated using a 20 nucleotide (nt) guide sequence, specific for the second exon of the GALR2 gene, as proposed by [25,26] and cloned into px330 vector in fusion with the transactivating crRNA (tracrRNA). Molecules of sgRNA and *S. pyogenes* Cas9 mRNA were transcribed in vitro using MEGAshortscript T7 Transcription Kit (Invitrogen) and mMESSAGE mMACHINE T7 Transcription Kit (Invitrogen), respectively, and purified using MEGAclear Transcription Clean-Up kit (Invitrogen) [27]. A single-stranded oligodeoxynucleotide (ssODN) donor was designed to contain the corresponding nucleotide alteration encoding the W248L missense variant, plus a one-point mutation to create a *ScaI* restriction enzyme site in the exon 2, three silent mutations to avoid cleaving DNA after homologous recombination and one silent mutation at the PAM (Protospacer Adjacent Motif). Five additional silent mutations were also introduced between the *ScaI* site and the missense mutation to disrupt the sgRNA recognition sequence on the donor strand, thereby preventing Cas9-mediated re-cleavage of successfully edited alleles and enhancing HDR efficiency. ssODN (200 bases) was purchased from IDT DNA, designed to be complementary to the target strand of sgRNA and with symmetric homology arms. The sgRNA (2µg), protein EnGen® Spy Cas9 NLS (5µg, NEB), and ssODN (2µg) were electroporated into C57BL/6J zygotes by the Laboratory of Animal Handling and Experimentation (LMEA, LNBio, CNPEM). Electroporated embryos were implanted into pseudopregnant CB6F1 (BALB/c × C57BL/6J) foster mothers. Mice were genotyped by PCR amplification from tail genomic DNA using *GALR2* CRISPR-PCR forward primer (5’ CATCCGCTACCCGCTGCACT 3’) and *GALR2* CRISPR-PCR reverse primer (5’ TAGCAGGCCCGCGCAGATTT 3’), followed by screening of animals by T7 Endonuclease I assay (T7EI) and/or *ScaI* restriction enzyme digestion. For T7EI assay, 600 bp PCR amplicon was purified by PCR purification kit (Qiagen, Singapore) and submitted to the assay with the T7 Endonuclease I enzyme (New England Biolabs, Ipswich, MA, USA), which recognizes and cleaves non-perfectly matched DNA. Briefly, 200 ng of founder’s PCR fragment was denatured at 95 ◦C for 10 min, followed by cooling annealing at –2 ◦C/second until 85 ◦C, and slow cooling at −0.1 ◦C/second until 25 ◦C. After the formation of DNA heteroduplexes, they were incubated with 5 U of T7EI enzyme (New England Biolabs) at 37 ◦C for 30 min. Then, the reaction was resolved in a 1% agarose gel electrophoresis [28]. Purified PCR amplicons were subjected to *ScaI* digestion to screen for successful mutagenesis, with cleavage of the 600 bp fragment into ∼400 bp and ∼200 bp bands indicating the presence of the desired mutations (Figure 1B). The presence of the mutations was further confirmed by Sanger sequencing (Figure 1C). KI animals were first crossed with WT C57BL/6J mice. Heterozygous offspring were then intercrossed for three generations to obtain homozygous KI and WT littermates, which were subsequently used in all experiments.

**Figure 1.**
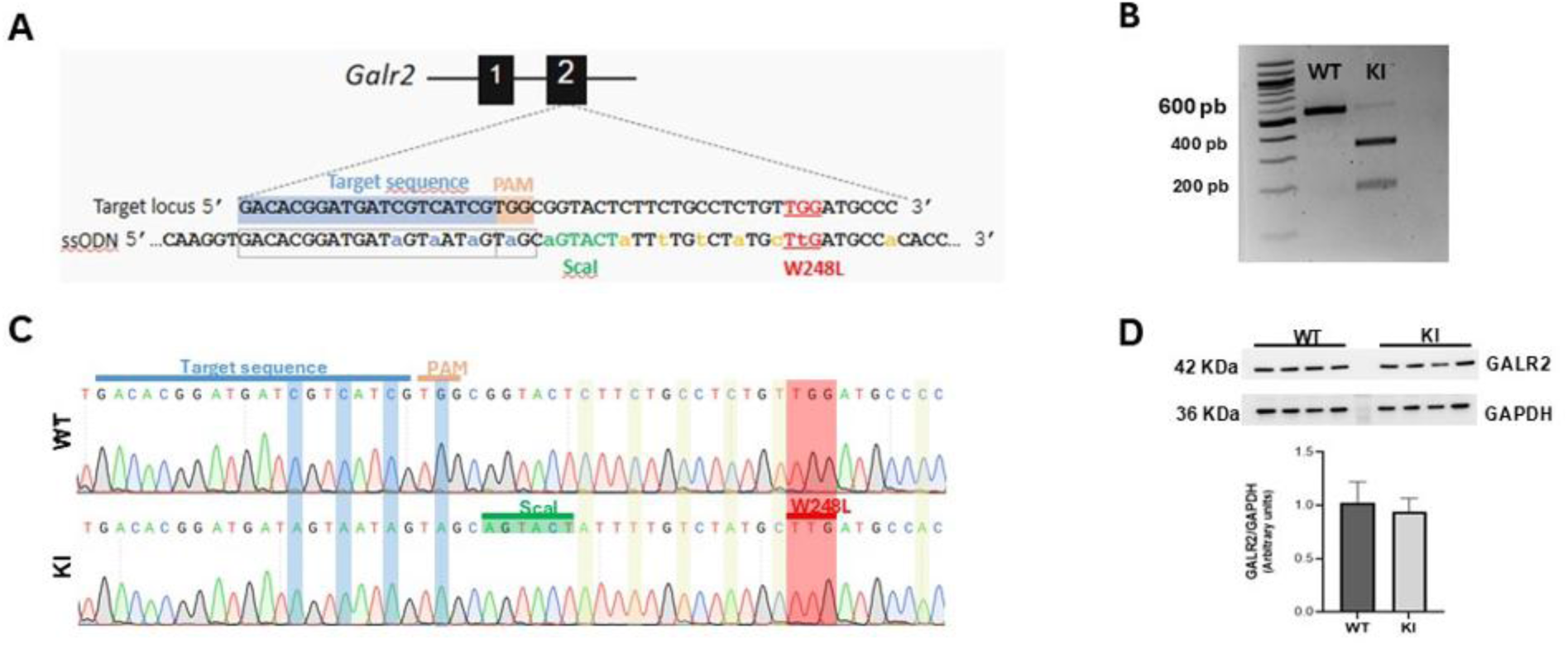
Generation and characterization of GALR2 W248L KI mice. **(A)** Schematic representation of the CRISPR/Cas9-mediated knock-in (KI) strategy targeting exon 2 of the *Galr2* gene, indicating the W248L missense mutation and silent substitutions introduced for ScaI-based genotyping and ssODN-mediated homologous recombination. **(B)** Representative agarose gel electrophoresis of ScaI-digested PCR products showing genotype-specific banding patterns for wild-type (WT) and homozygous KI mice. **(C)** Representative Sanger sequencing chromatograms confirming the W248L substitution in KI mice compared with WT controls **(D)** Representative Western blot analysis of GALR2 protein expression in brain cortical tissue from WT and KI mice, with densitometric quantification normalized to GAPDH (*n* = 4 independent experiments per group).

### Animals (Mice)

Ten-week-old male KI and WT mice were housed in the pathogen-free animal facility at the Animal Handling and Experimentation Laboratory (LMEA), at Brazilian Biosciences National Laboratory (LNBio, Campinas, Brazil), Brazilian Center for Research in Energy and Materials (CNPEM, Campinas, Brazil). Mice were housed in groups of 3–5 per microisolator cage under a 12:12 h light/dark cycle at 21–24 °C, with *ad libitum* access to food and water. All experimental procedures were conducted in accordance with the recommendations of the National Council for the Control of Animal Experimentation (CONCEA) and approved by the Ethics Committee on the Use of Animals (CEUA/CNPEM), under protocol #95. All efforts were made to minimize suffering.

### Induction and assessment of EAE

EAE was induced by subcutaneously immunizing mice in the flanks with MOG_35– 55_ (Proteimax Biotecnologia LTDA, Brazil). The 200 µg of administered MOG_35–55_ was composed of 100 µl PBS and 100 µl CFA (Sigma-Aldrich) supplemented with 5 mg/ml heat-inactivated *Mycobacterium tuberculosis* H37Ra (Difco). Additionally, mice received 200 ng pertussis toxin (Sigma-Aldrich) i.p. followed on the day of immunization as well as 2 days later. Clinical signs of EAE were scored on a standard 0–9 scale, according to previous recommendations [29], as follows: 0 = unaffected; 1 = partial limp tail; 2 = paralyzed tail; 3 = loss of coordinated movements and hindlimb paresis; 4 = one hindlimb paralyzed; 5 = both hindlimbs paralyzed; 6 = hindlimbs paralyzed and weakness in forelimbs; 7 = hindlimbs paralyzed and one forelimb paralyzed; 8 = hindlimbs and forelimbs paralyzed; and 9= moribund/death.

### Spinal cord Histology and Immunofluorescence

EAE-induced and sham mice were deeply anesthetized and transcardially perfused with ice-cold PBS followed by 4% paraformaldehyde (PFA). Spinal cords were collected and post-fixed in 4% PFA for 24 h. For histological analysis, tissues were dehydrated, paraffin-embedded, sectioned at 10 µm, and stained with hematoxylin–eosin or Luxol Fast Blue using standard protocols. Images were acquired using a Leica FS DM6 microscope at 10X magnification. For immunofluorescence, spinal cords were post-fixed in 4% PFA, cryoprotected in graded sucrose solutions (10%, 20%, and 30%), embedded in OCT, and cryosectioned at 12 µm. Sections were blocked with 1% BSA and incubated overnight at 4 °C with primary antibodies in PB containing 1% BSA and 0.2% Triton X-100. After washing, sections were incubated with species-specific secondary antibodies and with FluoroMyelin Red (1:1000; Invitrogen). Slides were mounted in glycerol-based medium. Primary antibodies included anti-Iba-1 (1:750), anti-GFAP (1:750), and anti-synaptophysin (1:1000). Histopathological evaluation of inflammatory infiltration and demyelination was performed in a blinded manner by a certified pathologist.

### Isolation of Leukocytes from the central nervous system (CNS)

EAE-induced and sham mice were deeply anesthetized and transcardially perfused with ice-cold PBS. The spinal cord was collected and minced with a sharp razor blade, following digestion for 30 min at 37°C with collagenase D (2.5 mg/ml; Roche Diagnostics). Mononuclear cells were isolated by the passage of the tissue through a cell strainer (70 µm), followed by centrifugation through a 37/70% Percoll gradient (GE Healthcare). For intracellular cytokine staining, isolated cells were stimulated as previously described, followed by flow cytometric analysis.

### Flow cytometry

CNS-isolated cells were washed and stained for 10 min at room temperature with fixable viability dye (Invitrogen) for dead cells exclusion and fluorochrome-labeled monoclonal antibodies against surface cell markers. For intracellular cytokine staining, cells were previously stimulated in culture medium with PMA (50 ng/ml; Sigma-Aldrich) and ionomycin (500 ng/ml; Sigma-Aldrich) for 4 h in the presence of monensin (GolgiStop 1.5 µg/ml; BD Biosciences) at 37°C in a humidified 5% CO_2_ chamber. Next, cells were fixed and permeabilized using the Foxp3 Transcription Factor Staining Kit (Invitrogen), followed by intracellular staining with monoclonal antibodies for 20 min. Data were acquired on a FACSVerse or machine (BD Biosciences) and analyzed using FlowJo software.

### Cytokine measurement

Supernatants from cell cultures were collected after centrifugation and IL-1β, IL-6, IL-10, IL-12/IL23 p40, CCL17 and CCL22 levels were measured by ELISA according to the manufacturer’s instructions (R&D Systems).

### Culture of Primary Hippocampal Neurons

Primary hippocampal neuronal cultures were prepared from postnatal day 0–1 (P0–1) C57BL/6J WT and GALR2 W248L mice of both sexes (8–12 neonates per group). Brains were dissected in ice-cold HBSS-based buffer, and hippocampi were isolated and enzymatically dissociated with 0.25% trypsin/EDTA for 20 min at 37°C. After enzymatic inactivation and DNase I treatment, tissues were mechanically dissociated, and cells were counted. Neurons were plated on poly-L-lysine-coated glass coverslips at ∼500 cells/mm² and maintained in neurobasal medium supplemented with B27, glutamine, and antibiotics. After 16 h, cultures were switched to maintenance medium containing 5 µM cytosine arabinoside (AraC). Cells were incubated at 37°C in 5% CO₂, with medium changes every 3– 4 days. Experiments were performed between 14 and 21 days in vitro (DIV14– 21) [30].

### Galanin Binding Assay in Primary Hippocampal Neurons

Human galanin (1–30; Santa Cruz Biotechnology) was conjugated in-house to the Cy3.5 fluorophore (Abcam) and purified by reverse-phase chromatography using an FPLC system. Labeled fractions were collected, solvent-evaporated, resuspended in Milli-Q water, and stored at 4 °C protected from light. Primary hippocampal neurons were plated at 1 x 10⁵ cells per well and maintained under standard culture conditions. Receptor binding assays were performed by live-cell confocal microscopy (Leica TCS SP8, 63x water-immersion objective) at 37 °C. Baseline images were acquired prior to ligand addition, followed by incubation with galanin-Cy3.5 (150 nM). Receptor binding was monitored for 20 min, with images acquired every 5 min. Image processing and quantitative analyses were performed using ImageJ software.

### Bone Marrow–Derived Macrophage Culture

Naïve mice were euthanized by ketamine/xylazine overdose, and femurs were aseptically collected. Bone marrow cells were obtained by flushing femoral cavities with RPMI-1640, centrifuged, and differentiated into bone marrow– derived macrophages (BMDMs) in LCCM-conditioned medium supplemented with M-CSF, fetal bovine serum, L-glutamine, and antibiotics. Cells were cultured in non-treated Petri dishes at 37 °C and 5% CO₂, with medium renewal on day 3. On day 6, macrophages were detached, collected, and resuspended in complete RPMI. Cells were counted, replated for ELISA (2 x 10⁵ cells/well) and flow cytometry (1 x 10⁶ cells/well), and allowed to rest for 2 h prior to experimental treatments.

### Bone Marrow–Derived Macrophage Polarization Assays

For polarization assays, bone marrow–derived macrophages were stimulated with lipopolysaccharide (LPS; 30 or 100 ng/mL; Sigma-Aldrich, St. Louis, MO, USA) for 24 h to induce M1 polarization, or with recombinant murine interleukin-4 (IL-4; 3 or 10 ng/mL; R&D Systems, Minneapolis, MN, USA) for 48 h to induce M2 polarization. Control cells were maintained in a complete RPMI medium. Cultures were incubated at 37 °C in a humidified atmosphere containing 5% CO₂. After stimulation, culture supernatants were collected and stored at −20 °C for cytokine quantification, and cells were harvested for flow cytometry analysis.

### NLRP3 Inflammasome Activation Assays

For NLRP3 inflammasome activation assays, bone marrow–derived macrophages were primed with lipopolysaccharide (LPS; 30 or 100 ng/mL; Sigma-Aldrich) for 4 h at 37 °C in a humidified atmosphere containing 5% CO₂, except for negative controls. After priming, the culture medium was replaced with a complete RPMI medium. To activate the NLRP3 inflammasome, cells were treated with nigericin (10 µM; Invitrogen) or ATP (5 mM; Sigma-Aldrich) for 1 h, except for negative controls. Subsequently, culture supernatants were collected and used for IL-1β quantification.

### Immunoblot analysis

Whole-cell lysates from cerebral cortex and hippocampus were prepared in RIPA buffer supplemented with protease and phosphatase inhibitors. Protein concentrations were determined by BCA assay. Equal amounts of protein were separated by SDS–PAGE and transferred to nitrocellulose membranes. Membranes were blocked with 5% nonfat milk in TBST and incubated overnight at 4 °C with anti-GALR2 primary antibody (1:1000; Abcam), followed by HRP-conjugated secondary antibody (1:5000). Immunoreactive bands were detected using ECL reagent and visualized with a ChemiDoc XRS system. Densitometric analyses were performed with ImageJ, and GAPDH was used as a loading control.

### Statistical analysis

GraphPad Prism 7.0 software was used for statistical analysis. Multiple-group comparisons were performed with either one way ANOVA or two-way ANOVA followed by Tukey’s post hoc test. Unpaired two-tailed Student’s t test was used for comparison of two conditions. Data are expressed as means ± SEM. P value < 0.05 was considered significant.

## Results

### Generation and functional validation of the GALR2 W248L knock-in mouse model

To investigate the biological relevance of W248L mutation in GALR2, we generated a knock-in mouse model in which tryptophan at position 248 was substituted by leucine (c.743G>C; p.248W>L) using CRISPR/Cas9-mediated homologous recombination (Fig. 1A). Correct targeting of the *Galr2 locus* was validated by PCR genotyping combined with restriction enzyme digestion and confirmed by Sanger sequencing (Fig. 1B-C). Homozygous knock-in (KI) mice exhibited normal viability and fertility with no overt developmental or behavioral abnormalities compared with WT littermates (Supplementary Fig. 1A-B). Furthermore, the GALR2 W248L mutation did not alter protein expression levels in the CNS (Fig. 1D), indicating that the mutation primarily affects receptor function rather than its expression.

After the demonstration by Garcia-Rosa et al., 2019 [24] that this substitution impairs GALR2 localization to the plasma membrane, inducing agonist-independent receptor internalization into endosomes, we exposed primary hippocampal neurons collected from WT and KI animals to fluorescently labeled galanin (galanin-Cy3.5) and monitored them by live imaging for 20 min. In WT neurons, galanin-Cy3.5 binding was readily detected along the plasma membrane as early as 5 min after incubation, with fluorescence intensity increasing over time. In contrast, KI neurons displayed a markedly reduced and barely detectable fluorescent signal throughout the imaging period (Supplementary Fig. 1C). Given that GALR2 protein levels in primary hippocampal neurons were comparable between genotypes (Supplementary Fig. 1D), the W248L mutation markedly reduced detectable galanin-Cy3.5 binding at the neuronal plasma membrane. This finding likely reflects reduced GALR2 availability at the cell surface, as previously observed [24].

### GALR2 W248L knock-in mice develop an exacerbated clinical course of EAE

To assess the impact of the GALR2 W248L mutation on neuroinflammation, homozygous KI and WT mice were immunized with MOG_35–55_ peptide to induce experimental autoimmune encephalomyelitis (EAE) (Fig. 2A). Disease course was monitored daily by clinical scoring and body weight measurements (Fig. 2B-C). The first neurological signs, characterized by tail weakness, appeared around day 11 post-immunization in both genotypes. Although disease onset was similar between groups, the subsequent trajectory and severity distribution differ notably. (Fig. 2C-D).

**Figure 2.**
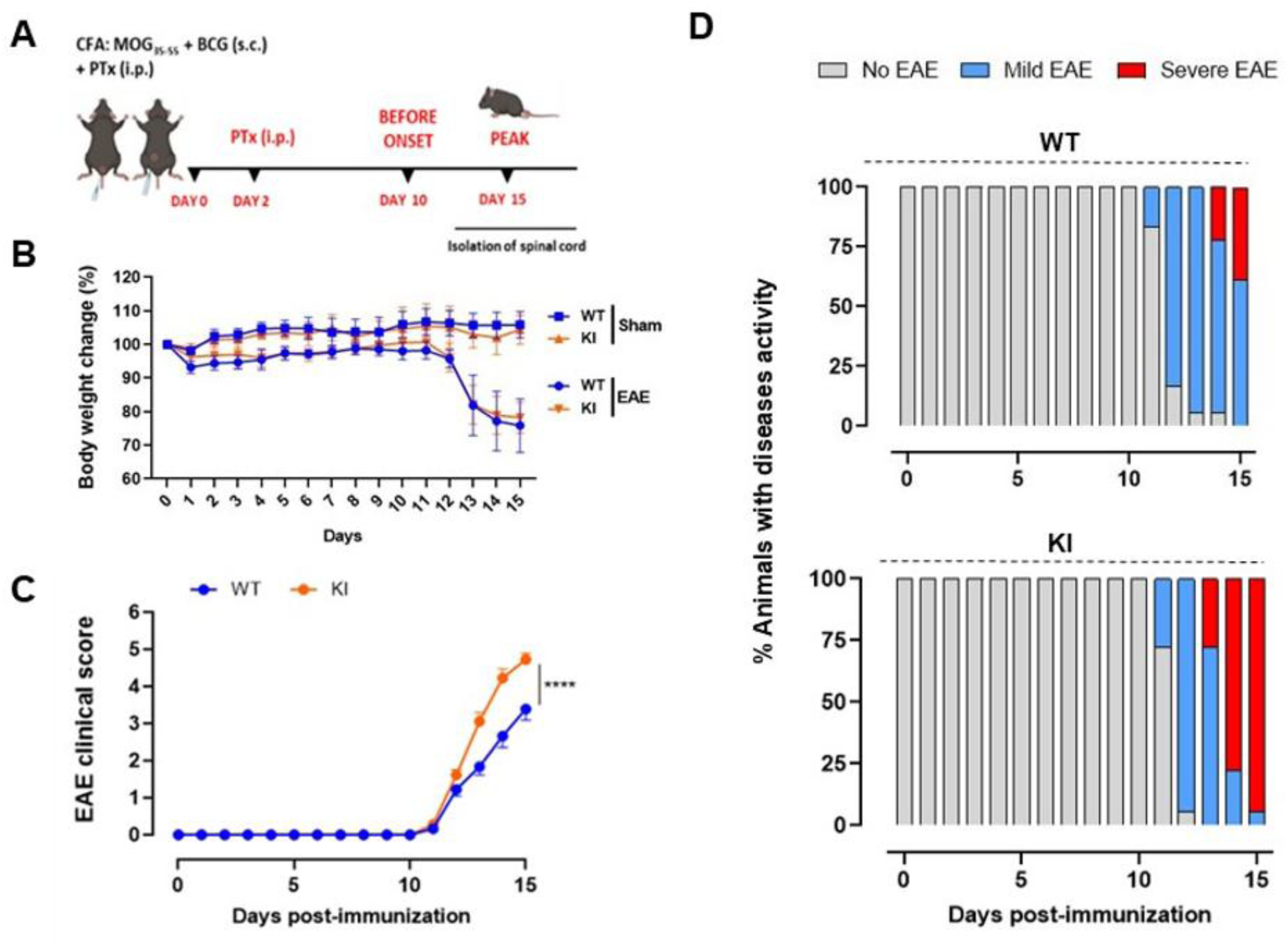
EAE progression and body weight changes in WT and GALR2 W248L mice. **(A-C)** EAE was induced in WT and GALR2 W248L Knock-in C57BL/6 mice by subcutaneous immunization with MOG_35-55_ and evaluated daily for body weight change and clinical signs (n = 18 per group). **(B)** Body weight variation throughout the days after immunization. **(C)** Cumulative EAE clinical scores. **(D)** Disease incidence by severity is represented on a bar chart as no EAE (score 0), mild EAE (score 1-3) and severe EAE (score 4-6). Data shown are representative of three independent experiments. Errors bars represent mean ± SEM. *P* values were determined by two-way ANOVA followed by Tukey’s post-hoc test. \*\*\*\**p* < 0.0001.

KI mice exhibited higher clinical scores throughout the disease course, with significantly elevated mean scores at the peak disease (day 15) compared with WT controls. Analysis of disease severity distribution revealed that KI mice progressed more rapidly from mild (score 1–3) to severe EAE (score 4–6) as compared to WT (Fig. 2D). By day 13, 27.7% of KI mice already exhibited unilateral hind limb paralysis, whereas this phenotype appeared in WT animals only one day later. On day 14, disease had progressed to severe stages in 77.7% of KI mice, compared with 22.2% of WT. At the experimental endpoint, severe EAE was observed in 94.4% of KI animals, in contrast to 38.8% of WT, and the maximum clinical score (6) was exclusively reached by KI mice (11.1%; *p* = 0.001). Together, these findings indicate that the GALR2 W248L mutation exacerbates EAE progression, suggesting that impaired GALR2 signaling may potentiate neuroinflammatory responses with implications for demyelinating pathology.

### Exacerbated EAE-induced spinal cord pathology and demyelination in GALR2 W248L Knock-in mice

Histopathological evaluation of the lumbar spinal cord confirmed the aggravated clinical course observed in GALR2 W248L KI mice. H&E staining revealed markedly increased mononuclear inflammatory infiltrates in KI compared with WT animals, with an inflammatory infiltrate that breached its usual confinement to the white matter, extending into the gray matter and diffusely invading the spinal cord parenchyma (Fig. 3A middle and lower right panels). FluoroMyelin staining provided higher spatial resolution of myelin disruption. Both genotypes exhibited focal loss of fluorescence in the white matter following EAE induction; however, KI mice showed broader depigmented regions and an overall disrupted white matter architecture. Strikingly, inflammatory infiltrates extended into the gray matter predominantly in KI animals, indicating a broader parenchymal involvement compared with WT counterparts (Fig. 3B lower right panels). Complementary Luxol Fast Blue (LFB) histochemistry showed preserved myelin in sham groups, whereas EAE-induced animals of both genotypes exhibited demyelinated regions characterized by LFB-negative areas, frequently associated with mononuclear infiltrates. In WT mice, demyelination was confined to small, discrete foci in the ventral white matter. By contrast, KI mice displayed a more diffuse and extensive demyelination pattern, with large regions of myelin loss often localized near blood vessels, a distribution reminiscent of perivascular pathology in progressive MS (Fig. 3C lower right). Taking together, these results indicate that the GALR2 W248L mutation exacerbates neuroinflammation and promotes more extensive demyelination in EAE.

**Figure 3.**
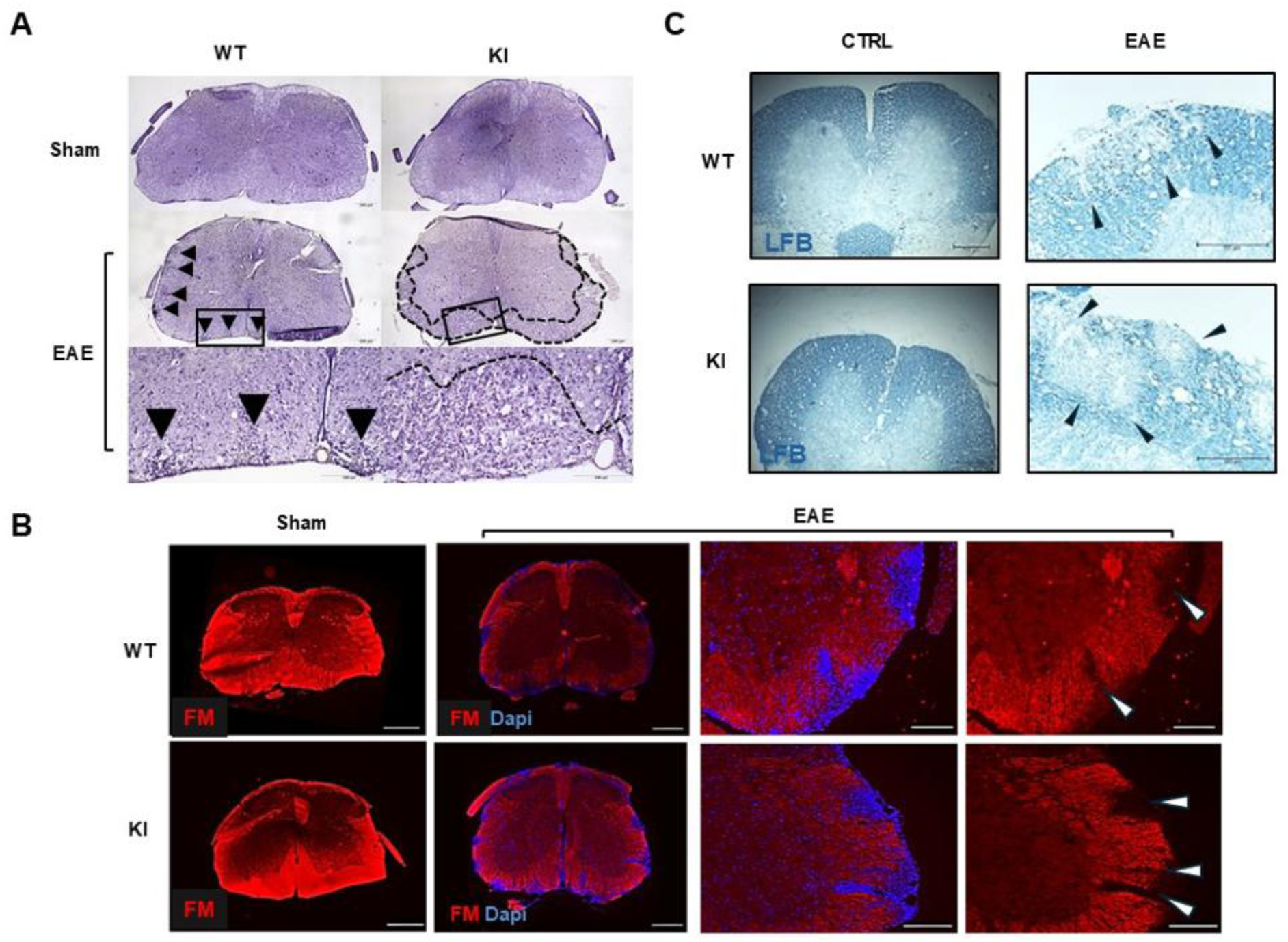
Exacerbated EAE-induced spinal cord pathology and demyelination in GALR2 W248L knock-in mice. **(A–C)** Transverse lumbar spinal cord sections from WT and GALR2 W248L knock-in (KI) mice under sham or EAE conditions, collected 15 days post-induction (sham n = 3; EAE n = 4 per group). **(A)** Inflammatory infiltration assessed by H&E staining at low and high magnification. WT-EAE mice show discrete subpial inflammatory foci (middle and lower left panels; arrowheads), whereas KI-EAE mice display a dense, band-like inflammatory infiltrate affecting the subpial white matter with extension into the gray matter (middle and lower right panels; dashed line). Scale bar: 200 µm. **(B– C)** Demyelination assessed by complementary histological approaches. **(B)** FluoroMyelin labeling (red) reveals focal myelin loss (upper and lower right panels; white arrowheads); nuclei are counterstained with DAPI (blue). Scale bar: 50 µm. **(C)** Luxol Fast Blue staining shows reduced myelin content, with demyelinated areas identified by attenuated staining at higher magnification (upper and lower right panels; black arrowheads). Scale bar: 200 µm.

### Flow cytometry analysis of lymphoid and myeloid CNS infiltrates

As part of the classical experimental workflow for the evaluation of EAE, and following the histopathological analyses of the spinal cord, we next investigated the immune cell profile within the CNS to explore potential mechanisms underlying the exacerbated clinical phenotype observed in KI mice. Flow cytometry analyses were performed on lymphoid and myeloid populations isolated from the spinal cord at the time of tissue collection, with corresponding data presented in Supplementary Fig. 2. Clinical monitoring up to 15 days post-induction (dpi) confirmed the more severe disease progression in KI mice compared with WT animals. However, flow cytometry analysis of spinal cord infiltrates revealed no significant differences in the proportion of CD4+ T cell populations (Th17/Treg) between WT and KI groups (Supplementary Fig. 2A-C). We also assessed the composition of the myeloid population in the spinal cord of EAE-induced animals. Likewise, the relative distribution of the myeloid compartment, including macrophages, neutrophils and dendritic cells did not differ significantly between KI and WT mice (Supplementary Fig. 2D-G). Collectively, these data reveal a complex, multifactorial immunological landscape in which the exacerbated clinical severity in KI mice cannot be explained by immune cell composition alone but instead implicates additional mechanisms in disease pathogenesis.

### Elevated microglial reactivity and pro-inflammatory shift in GALR2 W248L knock-in mice with EAE

To further characterize the neuroinflammatory response, we examined glial activation in the spinal cords of EAE-induced WT and KI mice at peak disease. Immunofluorescence staining for Iba1 revealed a marked increase in microglial activation in EAE-induced groups compared with sham controls, with KI mice displaying significantly greater microgliosis than WT animals (Fig. 4A-B). Morphological classification of microglia into surveillant (types I-II) and activated (types III-V) phenotypes demonstrated a pronounced shift toward activated morphologies in EAE-induced animals, particularly within Rexed lamina IX of the ventral gray matter (Fig. 4C). In sham controls, microglia predominantly exhibited a surveillant phenotype, characterized by small somata and highly ramified processes (Fig. 4D-F). In contrast, EAE induction significantly reduced the proportion of surveillant cells to 39.4%±3.35 (*p*<0.0001) in WT and 21.6%±3.07 in KI mice (p<0.0001), with activated morphologies predominating in KI animals (78.4%±3.07, p<0.0001) compared with WT (60.6%±3.35, p<0.0001).

**Figure 4.**
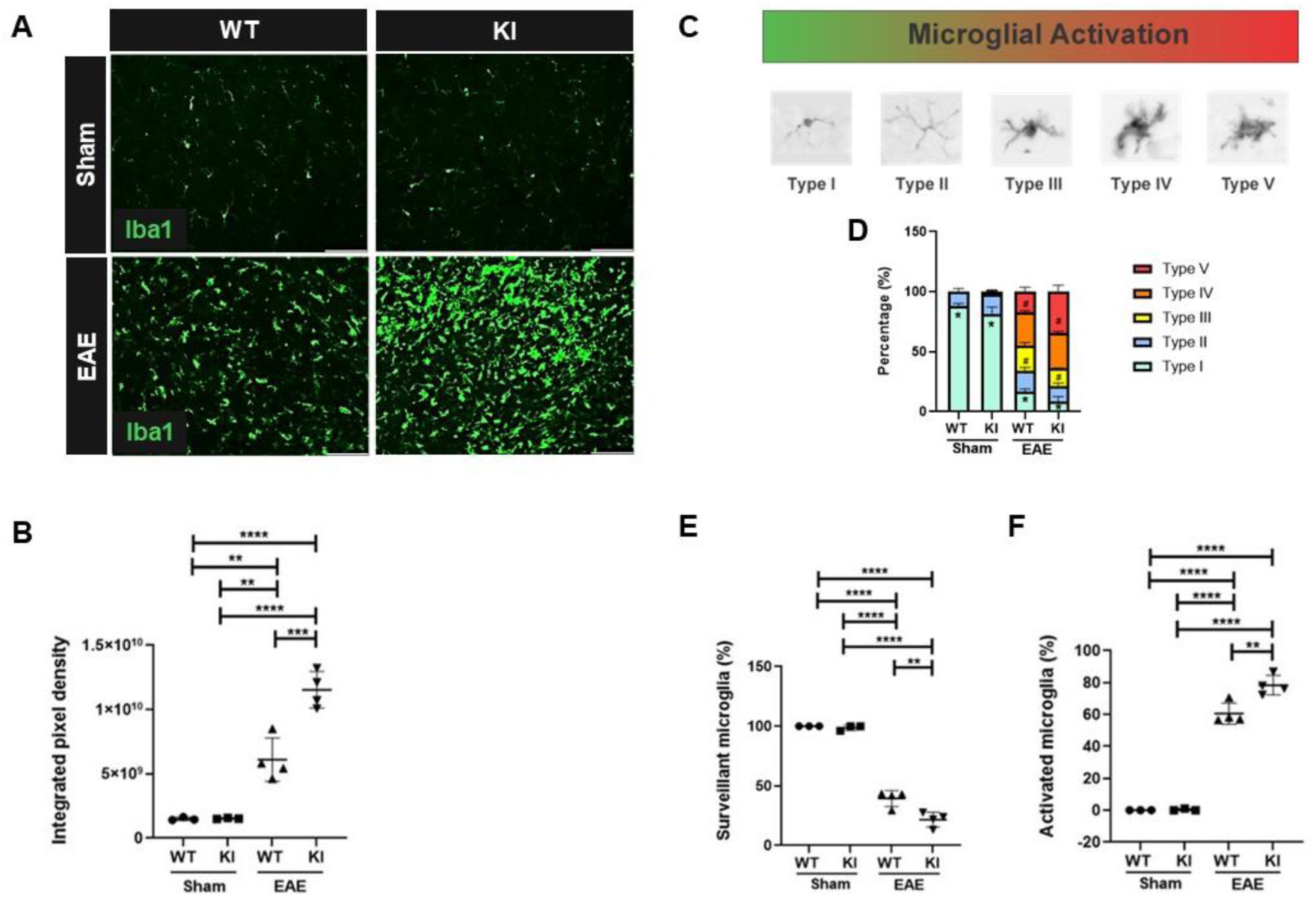
GALR2 W248L mutation promotes enhanced microglial reactivity and pro-inflammatory morphological activation during EAE. **(A)** Representative immunofluorescence images of the lumbar enlargement of spinal cord (L4–L6) immunolabeled with anti-Iba1 in WT and GALR2 W248L knock-in (KI) mice under sham or EAE conditions, collected at day 15 post-induction (sham n=3 and EAE n=4 per group). Scales bar represents 25 µm. **(B)** Quantification of microglial immunoreactivity expressed as integrated density of pixels (IDP) from anti-Iba1 immunolabeled images. Iba1 immunoreactivity was increased following EAE induction in both WT and KI mice compared with respective sham controls (WT, *p* = 0.0026; KI, *p* < 0.0001), with a significant difference between WT EAE and KI EAE groups (*p* = 0.0004). **(C)** Schematic representation of the microglial morphological categories used for classification, distinguishing surveillant (Type I and II) from activated microglia (Type III, IV and V). Scale bar indicates 10 µm. **(D)** Stacked-bar plots showing the relative distribution of microglial morphological subtypes (Types I–V) within Rexed lamina IX (ventral horn) across experimental groups. *P* values were determined by two-way ANOVA followed by Tukey’s post-hoc test. **(E)** Quantification of surveillant microglia (Types I–II). The proportion of surveillant microglia (Types I–II) was reduced following EAE induction in both genotypes (WT, *p* < 0.0001; KI, *p* = 0.0001), with differences between WT and KI groups under sham (*p* = 0.0027) and EAE conditions (*p* = 0.0020). **(F)** Quantification of activated microglia (Types III–V). The proportion of activated microglia (Types III–V) was higher in EAE groups compared with respective sham controls (WT and KI, *p* < 0.0001), with a difference between WT EAE and KI EAE groups (*p* = 0.0024). Error bars represent mean ± SEM. *P* values were determined by one-way ANOVA followed by Tukey’s post-hoc test. (\*\**p* < 0.01; \*\*\*\**p* < 0.0001).

### Astroglial reactivity and synaptic integrity in spinal cord during EAE are not altered by the GALR2 W248L mutation

Astrocytic activation is a pathological feature of EAE that appears to be independent of the GALR2 genotype, as demonstrated by immunofluorescence analysis with anti-GFAP, which showed an increase in astrocytic reactivity in EAE-induced animals compared to their respective sham (Supplementary Fig. 3A-B), confirming the involvement of astrocytes in the inflammatory response characteristic of the EAE model. Importantly, no significant differences were observed between EAE-induced WT and KI mice, further supporting that this astrocytic response represents a common pathological feature of disease induction rather than a genotype-dependent effect. Given the recognized roles of astrocytes and microglia in maintaining synaptic homeostasis, we next investigated whether the increased gliosis observed during EAE was associated with synaptic alterations in spinal motor neurons. Quantitative analysis revealed a significant reduction in synaptic coverage of motor neurons in EAE-induced animals compared with sham, consistent with gliosis-associated synaptic loss (Supplementary Fig. 3D). This reduction was evident in both WT and KI mice, with no statistically significant differences observed between genotypes at the peak of disease (15 dpi). A similar pattern was detected when assessing global synaptic preservation within the lateral motor nucleus of Rexed lamina IX, where both EAE groups differed significantly from their respective controls, but not from each other (Supplementary Fig. 3E). Although genotype-dependent differences were not evident at this time point, the more pronounced microglial activation observed in GALR2 W248L KI mice raises the possibility that prolonged disease evolution could lead to exacerbated synaptic disruption in this genotype.

### GALR2 W248L mutation enhances M1 polarization, suppresses M2 phenotype, and increases NLRP3 inflammasome activation in BMDMs

Given the increased microglial activation observed in EAE-induced GALR2 W248L KI mice, we next asked whether W248L-mediated GALR2 dysfunction intrinsically alters macrophage inflammatory programs using bone marrow– derived macrophages (BMDMs) (Fig. 5). BMDM polarization profiling revealed a pronounced shift toward a pro-inflammatory phenotype in KI cells (Fig. 5A-F). Under M1-polarizing conditions, as supported by flow cytometry analysis, we observed an increased CD86 expression in KI macrophages following stimulation with 100 ng/mL LPS, while CD80 expression did not differ between genotypes (Fig. 5A-B). KI BMDMs secreted significantly higher levels of IL-1β compared with WT at the highest LPS concentration, accompanied by increased IL-6 production at 30 and 100 ng/mL LPS, whereas IL-10 levels remained unchanged (Fig. 5D-F). In addition, IL-12/IL-23p40 expression was significantly elevated in KI macrophages, reaching maximal induction already at 10 ng/mL LPS, indicating an enhanced sensitivity to pro-inflammatory stimuli (Fig. 5C).

**Figure 5.**
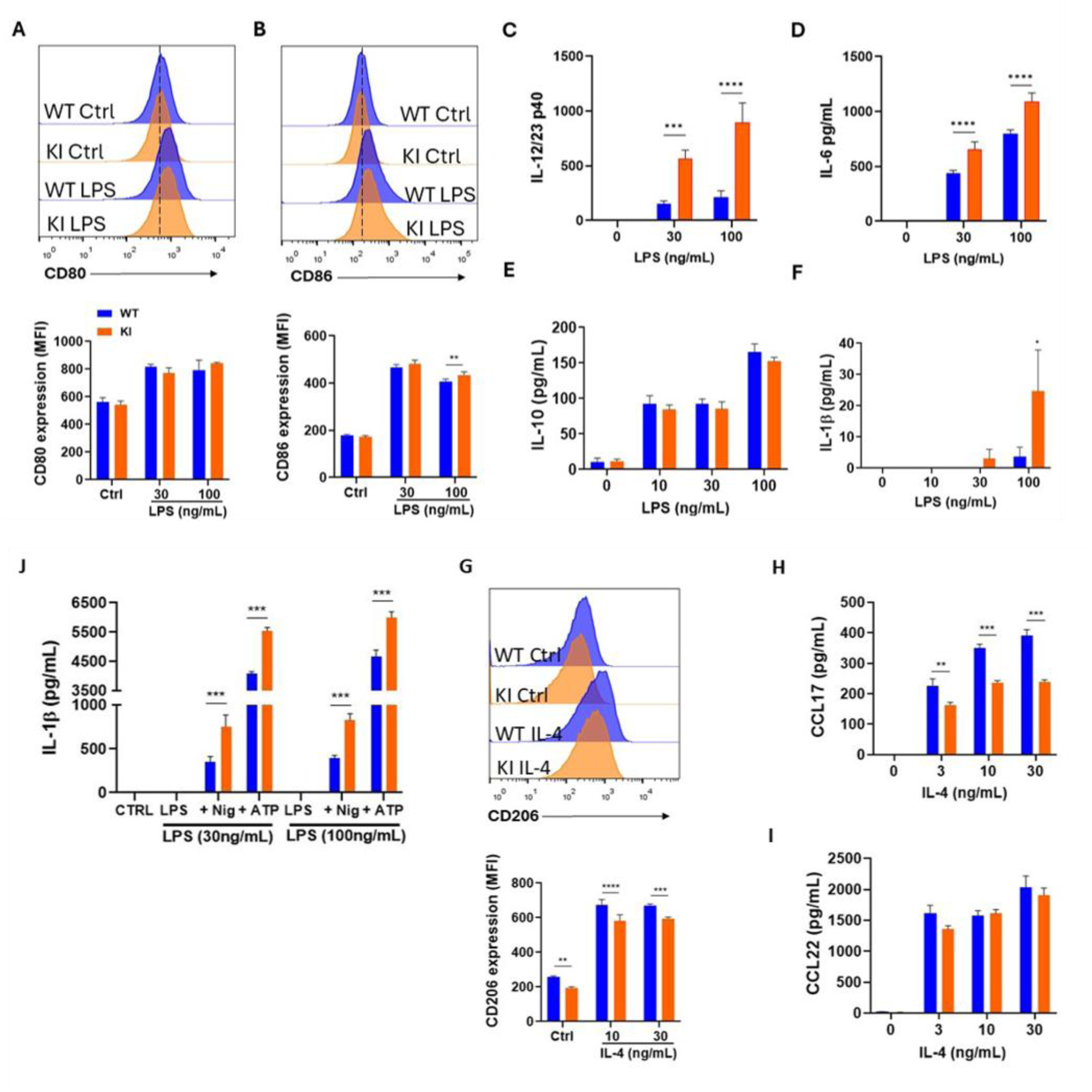
GALR2 W248L mutation promotes a pro-inflammatory macrophage profile through M1 enhancement, M2 impairment, and NLRP3 inflammasome activation. Bone marrow–derived macrophages (BMDMs) from naïve WT and GALR2 W248L knock-in (KI) mice were differentiated and stimulated as indicated. **(A–B)** After LPS stimulation (30 or 100 ng/mL) for 48 h, surface expression of the M1-associated markers **(A)** CD80 and **(B)** CD86 was quantified by flow cytometry. No differences were observed for CD80, whereas CD86 expression was increased in KI macrophages at 100 ng/mL LPS (*p* = 0.0071). **(C–F)** LPS-induced cytokine production was evaluated by ELISA after 24 h. **(C)** IL-12/IL-23 p40 levels were markedly increased in KI macrophages at 10, 30, and 100 ng/mL LPS (all *p* < 0.0001). **(D)** IL-6 was significantly elevated at 10 ng/mL (*p* = 0.0004) and 100 ng/mL LPS (*p* < 0.0001). **(E)** IL-10 secretion was not different between genotypes. **(F)** IL-1β levels were increased in KI macrophages at 100 ng/mL LPS (*p* = 0.0203). **(G–I)** For M2 polarization, BMDMs were stimulated with IL-4. **(G)** CD206 surface expression was significantly reduced in KI macrophages at 10 ng/mL (*p* < 0.0001) and 30 ng/mL IL-4 (*p* = 0.0002). **(I)** Secretion of the M2-associated chemokine CCL17 was decreased in KI macrophages at 3 ng/mL (*p* < 0.001) and at 10 and 30 ng/mL IL-4 (both *p* < 0.0001). **(I)** CCL22 levels were not significantly different between genotypes. **(J)** NLRP3 inflammasome activation was assessed following LPS priming (30 or 100 ng/mL) and subsequent stimulation with nigericin or ATP. IL-1β release was significantly increased in KI macrophages under all activation conditions tested (all *p* < 0.0001). Data are presented as mean ± SEM and are representative of three independent experiments. Statistical analyses were performed using two-way ANOVA followed by Sidák’s post-hoc test. (\*\**p* < 0.01; \*\*\*\**p* < 0.0001).

In contrast, M2 polarization assays revealed an impaired anti-inflammatory response in KI macrophages. Following IL-4 stimulation, flow cytometry analysis showed slightly reduced basal CD206 expression in KI cells. Moreover, although IL-4 induced CD206 upregulation in both genotypes, KI macrophages maintained significantly lower expression levels of CD206 (Fig. 5G). KI BMDMs exhibited a consistent reduction in CCL17 secretion across all tested concentrations, as well as decreased CCL22 levels at 3 ng/mL IL-4 (Fig. 5H-I). Together, these findings indicate that the GALR2 W248L mutation compromises the acquisition of an M2-like anti-inflammatory phenotype while favoring sustained M1-associated inflammatory responses.

Given the elevated IL-1β secretion in KI macrophages under M1-polarizing conditions, we next examined whether this phenotype was associated with enhanced NLRP3 inflammasome activation. BMDMs were stimulated with LPS alone or with LPS followed by Nigericin or ATP, and IL-1β release was quantified by ELISA. Under both LPS+Nigericin and LPS+ATP conditions, KI macrophages displayed significantly increased IL-1β secretion compared with WT controls (Fig. 5J), indicating enhanced NLRP3 inflammasome activation.

Collectively, these data establish that wild-type GALR2 signaling acts as a brake on macrophage inflammatory programs-restricting M1 polarization and inflammasome activity while favoring M2-related responses. The W248L mutation abrogates this protective regulation, tipping the balance toward a persistently pro-inflammatory macrophage phenotype, which may drive the heightened neuroinflammatory pathology seen in KI animals.

## Discussion

Multiple sclerosis (MS) is characterized by complex neuroimmune interactions involving glial activation, demyelination, and chronic inflammation. To determine whether impaired GALR2 signaling modifies neuroinflammatory responses in a model of MS-like pathology, this study provides the first *in vivo* functional characterization of the GALR2 W248L mutation using the experimental autoimmune encephalomyelitis (EAE) model. The data show that GALR2 W248L knock-in (KI) mice exhibit an earlier onset of serious neurological symptoms and a more severe clinical course of EAE compared with wild-type (WT) animals. These findings indicate that altered GALR2 functional signaling acts as a modifier of disease severity, amplifying neuroinflammatory responses, likely by weakening regulatory pathways that normally restrain innate immune activation.

Previous genetic studies identified the homozygous GALR2 W249L variant in a Brazilian MS patient [24], yet larger cohort studies failed to detect a clear association between GALR2 W249L or other variants in this gene [31] and MS susceptibility and disease severity. Although these observations argue against GALR2 mutations as primary drivers of MS pathogenesis, they do not exclude the involvement of molecular perturbations in other genes that converge on the same signaling axis. However, given the established roles of GALR2 in neuronal homeostasis, neurogenesis, neuronal integrity, and immune modulation, impaired galanin-GALR2 signaling pathway may act as a disease modifier that exacerbates neuroinflammation in the presence of additional genetic or environmental factors.

The clinical and histopathological profiles of the EAE-induced KI animals reinforce this interpretation. Spinal cord sections revealed extensive inflammatory infiltration and demyelination, with EAE-induced KI mice displaying diffuse invasion into the gray matter rather than the white-matter-restricted pattern observed in WT animals. This deeper parenchymal migration suggests enhanced inflammatory aggressiveness [32,33]. Quantitative flow cytometric analyses of spinal cord infiltrates revealed no significant genotype-dependent differences in the lymphoid or myeloid populations at the peak of EAE, indicating that quantitative changes in immune cell composition are unlikely to explain the aggravated phenotype in KI mice.

Immunohistochemical analyses revealed pronounced microglial activation in EAE-induced KI mice compared with WT. While astrocytic activation was comparable between genotypes, EAE-induced KI animals exhibited a marked increase in microglial density and a shift toward hypertrophic, ameboid morphologies, consistent with a sustained pro-inflammatory state [34]. These findings point to dysregulation of the innate immune system as a central contributor to disease exacerbation in the GALR2-mutant mice.

Both WT and KI exhibited reduced synaptic coverage of spinal motor neurons at the peak of disease, consistent with gliosis-associated synaptic remodeling in EAE [35]. Although no genotype-dependent differences were detected, the pronounced microglial activation in KI mice may suggest that prolonged disease evolution could result in exacerbated synaptic loss. Excessive microglial activation has been shown to create a toxic *milieu* for oligodendrocytes and neurons resulting in increased demyelination, synaptic destabilization and neurodegeneration, processes that are likely amplified when endogenous anti-inflammatory signaling is compromised [36,37].

In line with the prominent involvement of innate immune mechanisms, the intrinsic impact of the GALR2 W248L mutation on macrophage inflammatory programs was then examined. *Ex vivo* analyses revealed a clear shift in macrophage polarization toward a pro-inflammatory state in bone marrow-derived macrophages from KI mice. Specifically, KI macrophages exhibited enhanced M1-associated responses and a reduced capacity to acquire an M2-like phenotype. Moreover, elevated IL-1β secretion and increased NLRP3 inflammasome activation were observed in KI macrophages under inflammatory conditions. Together, these findings indicate that GALR2 signaling contributes to the maintenance of macrophage polarization balance and the restraint of inflammasome activation, whereas its disruption favors sustained pro-inflammatory programs.

At the mechanistic level, GALR2 signals mainly through G_q/11_ proteins, which activate phospholipase C, leading to IP₃ and DAG generation, intracellular calcium mobilization, and subsequent protein kinase C (PKC) activation [38]. Disruption of this signaling axis may impair calcium-dependent regulatory processes in innate immune cells that are critical for controlling inflammatory responses [21,39,40,41]. Attenuation of these pathways is therefore likely to bias immune cells toward persistent pro-inflammatory activation. In addition to canonical G protein–mediated signaling, emerging evidence indicates that GALR2 can engage β-arrestin–dependent pathways in a ligand- and context-specific manner. Notably, a recent study demonstrated that biased activation of GALR2 toward β-arrestin-2 signaling exerts potent anti-inflammatory effects, independently of Gq/11 coupling, by dampening inflammatory responses and preserving tissue integrity [42]. Thus, the GALR2 W248L mutation may not only reduce overall receptor availability at the plasma membrane but also favor a maladaptive signaling profile that promotes sustained innate immune activation.

Consistent with this model, live-cell imaging of primary hippocampal neurons revealed markedly reduced galanin binding in KI-derived cultures, despite preserved receptor expression. This observation aligns with previous reports showing that the W248L/W249L mutant receptor accumulates in endosomal compartments rather than localizing to the plasma membrane [24].

Reduced GALR2 surface availability is therefore expected to attenuate galanin-dependent neuroprotective and immunomodulatory signaling [14,39,43,44,45]. Accordingly, this diminished receptor accessibility may contribute to increased susceptibility to neuroinflammation and demyelination observed in EAE-induced KI animals.

In conclusion, this study provides the first experimental evidence that the GALR2 W248L substitution influences the course of neuroinflammation in experimental autoimmune encephalomyelitis. Impaired GALR2 signaling is associated with an earlier onset and greater severity of clinical signs, accompanied by intense inflammatory infiltration, exacerbated microglial activation, M1 macrophage predominance, enhanced NLRP3 inflammasome activity, and more extensive demyelination. While this impaired GALR2 variant alone is unlikely to initiate MS, it appears to act as a potent modifier of disease severity by weakening regulatory pathways that normally constrain neuroinflammation. These results highlight the galanin-GALR2 axis as a critical interface between neuronal and innate immune signaling and suggest that modulation of this pathway may represent a promising strategy for controlling neuroinflammation in demyelinating disorders.

## Statements

## Acknowledgements

This research used facilities of the Brazilian Biosciences National Laboratory (LNBio), part of the Brazilian Center for Research in Energy and Materials (CNPEM), a private non-profit organization under the supervision of the Brazilian Ministry for Science, Technology, and Innovations (MCTI). The staffs of Animal Handling and Experimentation Laboratory (LMEA; proposal 20230476), Laboratory of Bioimaging (LIB; proposal 20250514) and Mammalian and Insect Cell Culture Laboratory (LCCMI; proposal 20232701) facilities are acknowledged for assistance during the experiments.

## Authors contributions

A.S. conceived and designed the study, supervised the project, wrote, corrected, and critically revised the article.

D.B.B.T. contributed to study conceptualization and revised the article.

R.M.N. performed animal experiments and cellular assays, analyzed and interpreted the data, and wrote the article.

D.C.G. assisted with neuronal imaging experiments, contributed to conceptualization, and revised the article.

K.Y.D. provided technical support for the generation of the knock-in mouse lineage, assisted with animal experiments, performed Western blot experiments, and revised the article.

J.P.M.L. and M.S.B. assisted with macrophage polarization experiments, NLRP3 inflammasome activation assay and revised the article.

L.E.A.D. performed flow cytometry analyses, assisted with phenotyping of lymphoid and myeloid populations in the spinal cord, and revised the article.

S.J.O.P. assisted with immunofluorescence staining and imaging experiments and revised the article.

A.A.S. performed the histopathological evaluation and assisted with histopathological analyses.

E.D.N. initial discussions of the project and article revision.

A.L.R.O. provided facilities for immunohistochemistry experiments and contributed to data analysis and interpretation and revised the article.

J.C.A.F. provided facilities for flow cytometry, macrophage polarization, and inflammasome activation assays, and contributed to experimental design, data interpretation and revised the article.

All authors have reviewed the final version of the manuscript and approved its submission

## Ethics declarations

### Ethics approval

All procedures involving animals were approved by the Institutional Animal Care and Use Committee (Protocol No. 95) of Brazilian Center for Research in Energy and Materials (CNPEM).

## Abbreviations

AraC: Cytosine arabinoside
ATP: Adenosine triphosphate
BBB: Blood–brain barrier
BMDM: Bone marrow–derived macrophage
cAMP: Cyclic adenosine monophosphate
CFA: Complete Freund’s adjuvant
CNS: Central nervous system
CRISPR: Clustered regularly interspaced short palindromic repeats
DAG: Diacylglycerol
DAMP: Damage-associated molecular pattern
DIV: Days in vitro
dpi: Days post-immunization
EAE: Experimental autoimmune encephalomyelitis
ELISA: Enzyme-linked immunosorbent assay
ERK: Extracellular signal-regulated kinase
Foxp3: Forkhead box P3
GAPDH: Glyceraldehyde-3-phosphate dehydrogenase
GALR1: Galanin receptor type 1
GALR2: Galanin receptor type 2
GALR3: Galanin receptor type 3
GPCR: G protein-coupled receptor
H&E: Hematoxylin and eosin
IFN-γ: Interferon gamma
IL: Interleukin
IP₃: Inositol trisphosphate
KI: Knock-in
LFB: Luxol fast blue
LPS: Lipopolysaccharide
MAPK: Mitogen-activated protein kinase
MOG: Myelin oligodendrocyte glycoprotein
MS: Multiple sclerosis
NLRP3: NOD-like receptor family pyrin domain containing 3
OCT: Optimal cutting temperature compound
PAM: Protospacer adjacent motif
PKA: Protein kinase A
PKC: Protein kinase C
PLC: Phospholipase C
ROS: Reactive oxygen species
sgRNA: Single guide RNA
SNV: Single-nucleotide variant
ssODN: Single-stranded oligodeoxynucleotide
T7EI: T7 endonuclease I
Th17: T helper 17 cell
TNF-α: Tumor necrosis factor alpha
WT: Wild type

