## Supplementary Figures (1-3) for "GALR2 W248L mutation exacerbates neuroinflammation through pro-inflammatory macrophage polarization and microglial activation in experimental autoimmune encephalomyelitis"

**Affiliations:**

The supplementary information is a single file that includes:

- Three Supplementary Figures (1-3), including one that complements the main figure.

**SUPPLEMENTARY FIGURES AND LEGENDS**

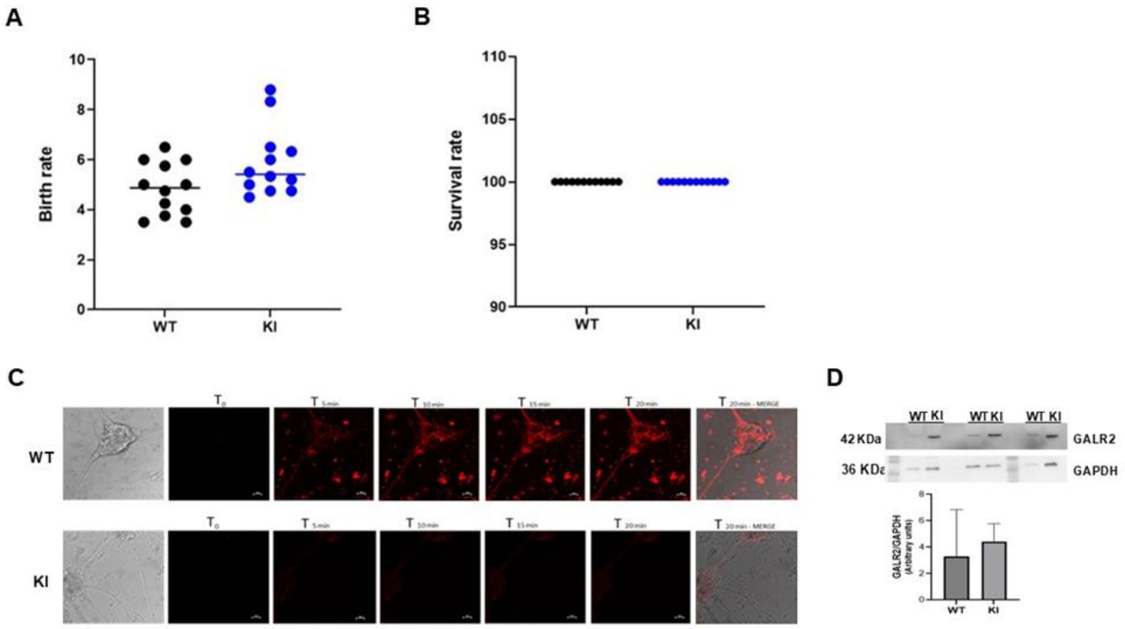


**Supplementary Figure 1 (related to figure 1). Reproductive performance, survival, and neuronal GALR2-related neuronal characterization in WT and GALR2 W248L mice. (A)** Birth rate expressed as the percentage of viable pups per litter in WT and KI mice (WT, $n$ = 18; KI, $n$ = 20 litters; unpaired $t$-test). **(B)** Survival analysis of WT and KI mice over the observation period (WT, $n$ = 287; KI, $n$ = 230 animals; log-rank test). **(C)** Representative live-cell confocal microscopy images of primary hippocampal neurons from WT and KI mice incubated with fluorescently labeled galanin (galanin–Cy3.5), acquired at baseline ($T_0$) and at the indicated time points (5–20 min) after ligand addition. **(D)** Representative Western blot analysis of GALR2 protein expression in protein lysates from cultured primary hippocampal neurons isolated from WT and KI mice, with densitometric quantification normalized to GAPDH ($n$ = 3 independent experiments per group).

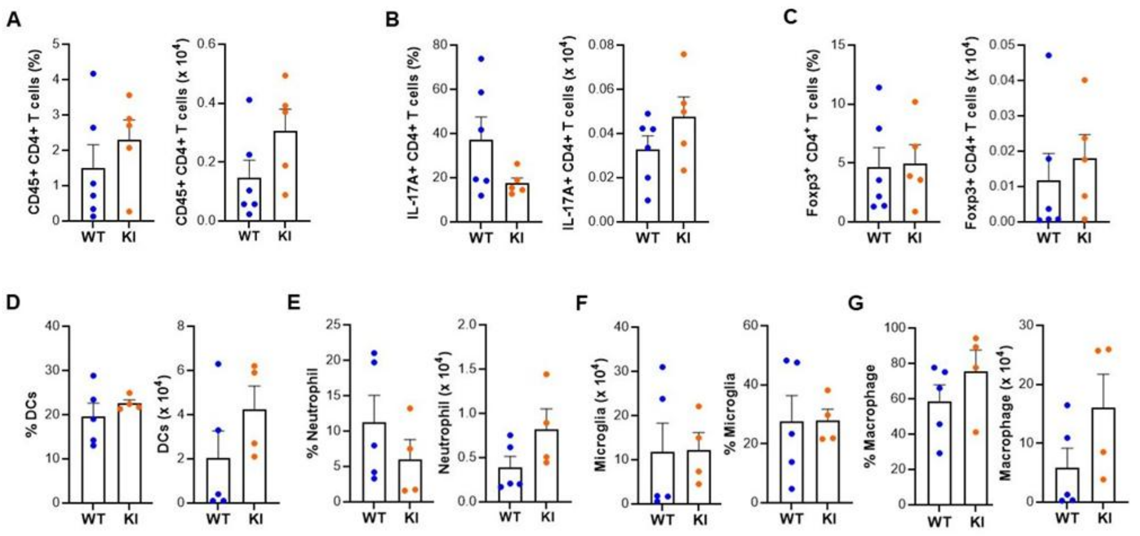


**Supplementary Figure 2. Comparative flow cytometric analysis of lymphoid and myeloid immune populations in WT and GALR2 W248L knock-in mice during EAE.** Spinal cord–infiltrating leukocytes were isolated from WT and GALR2 W248L knock-in (KI) mice 15 days after EAE induction and analyzed by flow cytometry. Data are presented as relative frequencies (%) and absolute cell numbers. **(A)** $CD45^{+}$ $CD4^{+}$ T cells. **(B)** IL-$17A^{+}$ $CD4^{+}$ T cells. **(C)** $CD4^{+}$ $Foxp3^{+}$ regulatory T cells. **(D)** Dendritic cells. **(E)** Neutrophils. (**F)** Microglial cells. **(G)** Infiltrating macrophages. Data shown are representative of two independent experiments for lymphoid analyses and three independent experiments for myeloid analyses. Error bars represent mean ± SEM. P values were determined using two-way ANOVA followed by Tukey's post-hoc test.

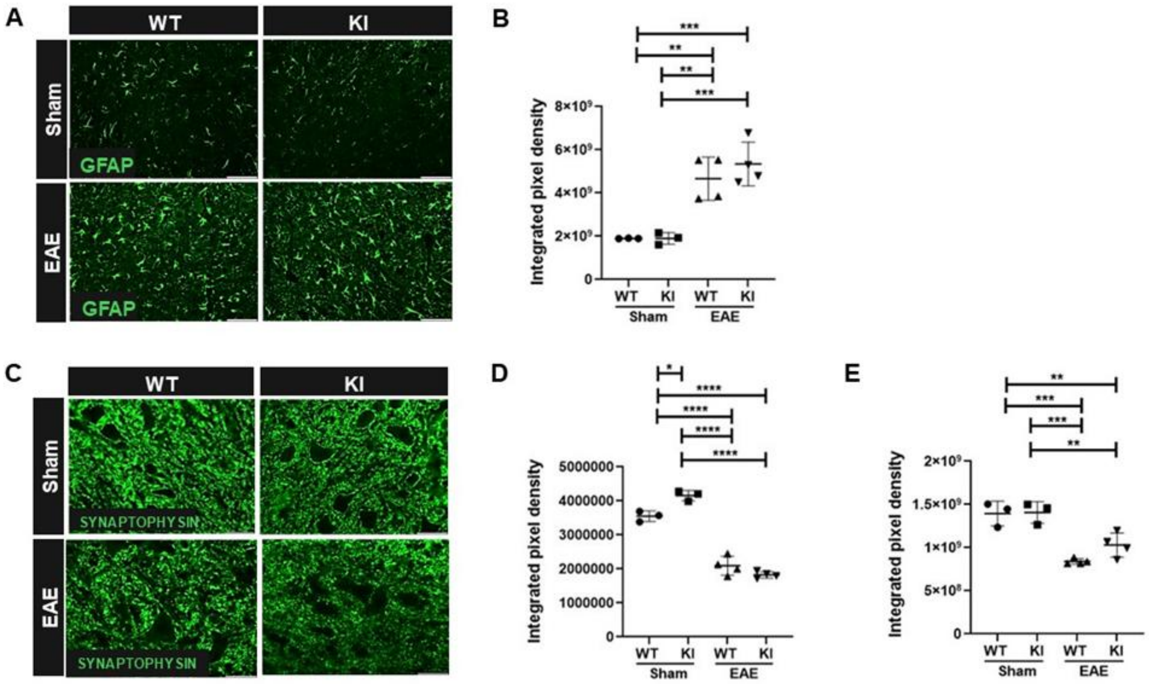


**Supplementary Figure 3. Astroglial reactivity and synaptic integrity in spinal cord during EAE are not altered by the GALR2 W248L mutation. (A and C)** Representative immunofluorescence images from the lumbar enlargement (L4–L6) of WT and GALR2 W248L knock-in (KI) mice under sham or EAE conditions, collected at day 15 post-induction (sham n = 3; EAE n = 4 per group). Scale bar: 25 µm. **(A)** Astrocytes immunolabeled with anti-GFAP. **(B)** Quantification of astrocytic GFAP immunoreactivity expressed as integrated density of pixels (IDP). **(C)** Synaptic terminals immunolabeled with anti-synaptophysin in Rexed lamina IX (ventral horn). **(D)** Quantification of synaptophysin immunoreactivity (IDP) associated with spinal motoneurons. **(E)** Quantification of synaptophysin immunoreactivity (IDP) in the lateral motor nucleus. GFAP immunoreactivity was increased following EAE induction in both WT and KI mice compared with respective sham controls. Synaptophysin immunoreactivity was reduced after EAE induction in both genotypes at the level of motoneurons and the lateral motor nucleus. No genotype-dependent differences were detected under EAE conditions. Data are presented as mean ± SEM. Statistical significance was determined using two-way ANOVA followed by Duncan’s multiple range test.
